# Molecular determinants and bottlenecks in the unbinding dynamics of biotin-streptavidin

**DOI:** 10.1101/176594

**Authors:** Pratyush Tiwary

## Abstract

Biotin-streptavidin is a very popular system used to gain insight into protein-ligand interactions. In its tetrameric form, it is well-known for its extremely long residence times, being one of the strongest known non-covalent interactions in nature, and is heavily used across the biotechnological industry. In this work we gain understanding into the molecular determinants and bottlenecks in the unbinding of the dimeric biotin-streptavidin system in its wild type and with N23A mutation. Using new enhanced sampling methods with full atomistic resolution, we reproduce the variation caused by N23A mutation in experimentally reported residence time. We also answer a longstanding question regarding cause/effect in the coupled events of bond stretching and bond hydration during unbinding and establish that in this system, it is the bond stretching and not hydration which forms the bottleneck in the early parts of the unbinding. We believe these calculations represent a step forward in the use of atomistic simulations to study pharmacodynamics. An improved understanding of biotin-streptavidin unbinding dynamics should also have direct benefits in biotechnological and nanobiotechnological applications.

---

The interaction of streptavidin with biotin in the homotetramer form is one of the strongest known non-covalent interactions in nature.^1–4^ This system forms a central component of countless biological, biotechnological and nano-biotechnological applications, and has long been used as a model system to understand the molecular basis of high affinity binding.^5,6^ Designing specific mutations in streptavidin allows tailoring a wide range of affinities and rate constants^7^ as needed for diverse practical applications. Further, given that many of the structural motifs found in biotin-streptavidin are utilized in numerous other protein-ligand interactions, it is very desirable to map out the structural and dynamical motifs underlying the slow dissociation kinetics in this system.

Experiments have reported both the binding affinity (*K*_*D*_) and the binding/unbinding rate constants (*k*_*on*_, *k*_*off*_ respectively, with *K*_*D*_ = *k*_*on*_/*k*_*off*_) for this system with and without mutations in the host. Of these numbers, the unbinding rate constant *k*_*off*_ (also often represented through its inverse, the residence time) has recently received a lot of attention as possibly being a much better predictor of drug efficacy compared to the static *K*_*D*_.^8–10^ While experiments can provide a direct measurement of *k*_*off*_, it is not so easy to derive from experiments direct information into the molecular determinants of the residence time. These determinants could come from a diverse range – protein-ligand contacts, solvation, protein/ligand flexibilities and many others. Having an atomistic resolution understanding of the role played by these determinants in the unbinding dynamics, one could then propose structural modifications that would assist in rational design of drugs with desired residence times. With the advent of petaflop computing and reliable force-fields, all-atom molecular dynamics (MD) simulations seem ideally poised to gain atomic resolution insight into the process of ligand unbinding. However, the reported residence times of most ligands of practical interest are well into the millisecond timescale and much slower, which is far beyond what can be attained with atomistic simulations even with the fastest super-computers.

As such, in this work, we use recent developments in enhanced sampling methods that complement atomistic simulations in a controllable manner.^11–13^ These give direct observations of kinetic information^14–20^ – not just the unbinding rate constant, which experiments can measure reliably, but also other entities which are very hard to obtain from experiments, such as unbinding pathways, intermediate states, their residence times, and the rate-determining steps.

Inspired by the pioneering work of Grubmüller, Heymann and Tavan,^5^ who used out-of-equilibrium pulling to study the unbinding of a biotin-streptavidin monomer, in this work we look at these two configurations in the symmetric dimer unit of biotin-streptavidin (Fig. 1) which we expect to be more tractable computationally, given that its binding affinity is around 6 orders of magnitude lower than for the full tetrameric system.^21^ With the new enhanced sampling methods^11,12^ we study the biotin-streptavidin system (PDB ID: 3ry2) in its wild-type (WT) and with an Asparagine-Alanine mutation (N23A).^22^ We expect and find that considering the dimer instead of tetramer brings down the residence time from more than a day to around a second, which is a timescale regime where the enhanced sampling methods of this work have been shown to be reliable.^14–16^ Note that there is no fundamental reason why these methods should not work for much longer timescales, and this will be the subject of future investigations. For the tetrameric biotin-streptavidin system the WT and N23A have been reported to have *k*_*off*_ of 4.4 ± 0.3 × 10^-5^*s*^-1^ and 1030 ± 220 × 10^-5^*s*^-1^ respectively.^22^ Thus, a single point mutation diminishes the residence time by a staggering nearly 250 times. Our simulations reproduce this behavior in the dimeric biotin-streptavidin and give crucial insight into why it happens. Our calculated residence times are also in rough order of magnitude range expected from the reported difference in binding affinity for the tetrameric versus dimeric systems.^21^

**FIG. 1:**
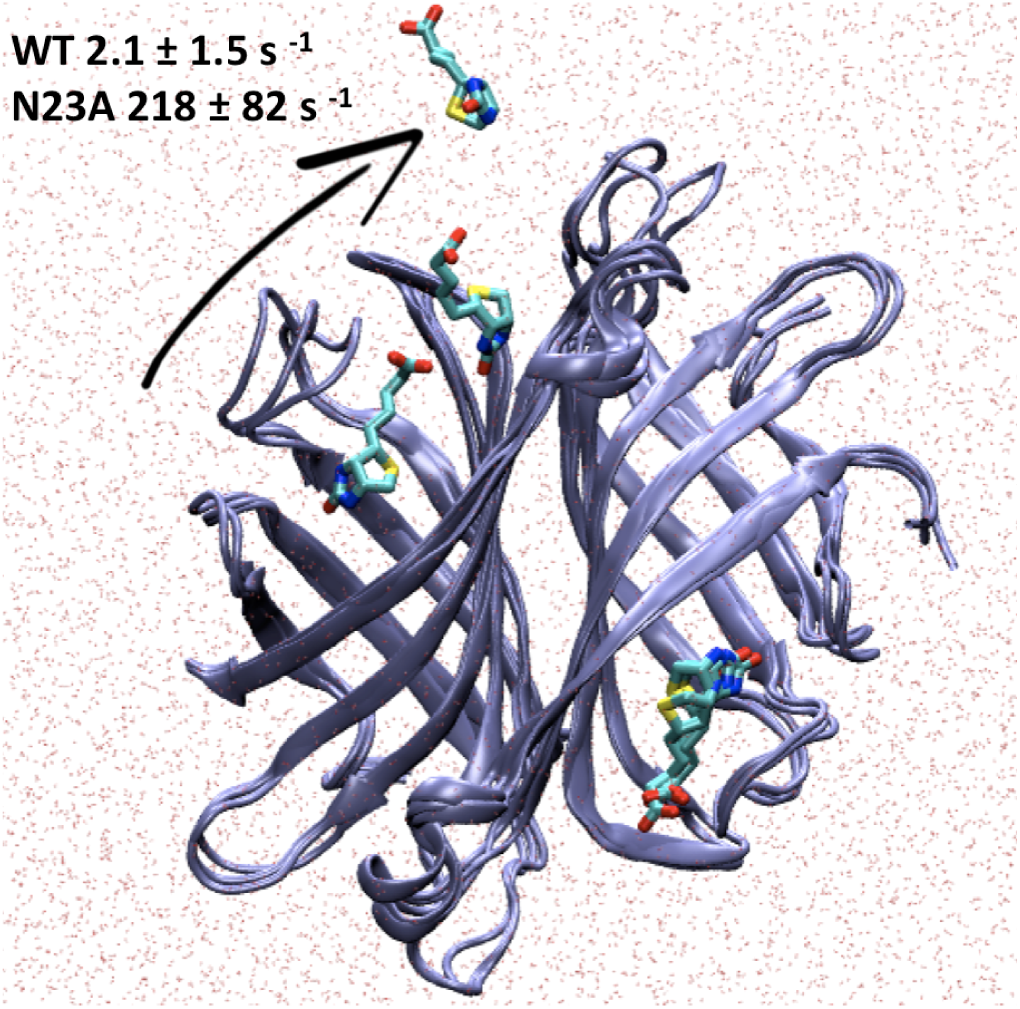
Symmetric dimeric unit of biotin-streptavidin in explicit water molecules, with superimposed snapshots of the host and ligand in bound, unbound and an intermediate state as obtained from molecular dynamics trajectories in this work. Note the presence of two units of biotin and streptavidin. Simulations in this work consider the unbinding in one of the two units. The off-rate *k*_*off*_ as found in this work (see Results) are also provided for WT and N23A.

Furthermore, we settle a long-standing question about the cause versus effect nature of two events, namely water movement into binding pocket and lengthening of a specific biotin-streptavidin hydrogen bonding interaction (D128-biotin bond). Previous simulations have demonstrated that these two events happen in a coupled manner before unbinding can proceed.^2,3^ However they could not comment on the relative importance of the two events. In this work, using the recently proposed method “Spectral gap optimization of order parameters (SGOOP)” proposed by Tiwary and Berne,^12^ we optimize the reaction coordinate (RC) for biotin unbinding which quantifies the relative importance of these two and other molecular determinants in unbinding. We establish that it is the hydrogen bond stretching that forms the bottleneck for the unbinding to proceed. Using the optimized RC for the WT and N23A forms, we then perform multiple independent “infrequent metadynamics” simulations.^11,13^ These give direct estimates of the various unbinding timescales and pathways. All kinetic observables are validated througbh statistical self-consistency tests.

## RESULTS

We start with a trial RC (Ψ_1_ in Table 1) for both WT and N23A forms, with which we perform preliminary metadynamics unbinding simulations to explore the free energy. Given the rough nature of the RC these runs are only for preliminary exploration of the configuration space. We also form short unbiased MD runs in the bound pose to calculate dynamical observables for the Maximum Caliber framework^23,24^ needed by SGOOP (Methods). These metadynamics and unbiased MD runs are input into SGOOP which processes them to optimize the RC as a linear combination of 5 order parameters (Table 1). Note that SGOOP as implemented in this work through the Maximum Caliber framework, takes into explicit account the diffusion anisotropy between different collective variables^25^ which has been known to have profound effects on reaction pathways as has been show for ligand binding and many other problems.^26–29^ A second set of “infrequent metadynamics” runs is performed using this optimized RC, with much slower bias deposition frequency in order to not perturb the transition states and meet the requirements of Ref.^11,30^ (Methods). 14 such independent runs each were performed for the WT and N23A forms. All sets of metadynamics runs (i.e. for input into SGOOP as well as infrequent metadynamics) were started in the bound x-ray pose and stopped when the ligand was fully unbound and freely diffusing. Both sets of unbinding times pass the Kolmogorov-Smirnoff test of Ref.^30^ as well as other statistical analyses reported in this section and in supplementary results. Our calculated *k*_*off*_ values are 2.1 ± 1.5 *s*^-1^ and 218 ± 82 *s*^-1^ respectively for the WT and N23A - thus we find the mutant to be around 100 times faster at unbinding than the wild type. Note that these are in rough agreement with the *k*_*off*_ values for the full tetrameric system reported in the introduction taking into account that the dimeric system has 5-6 order of magnitude smaller binding affinity *K*_*D*_, and that if one assumes *k*_*on*_ to be diffusion limited, most of the variation in *K*_*D*_ = *k*_*on*_*/k*_*off*_ should come from *k*_*of f*_.

**TABLE I:**
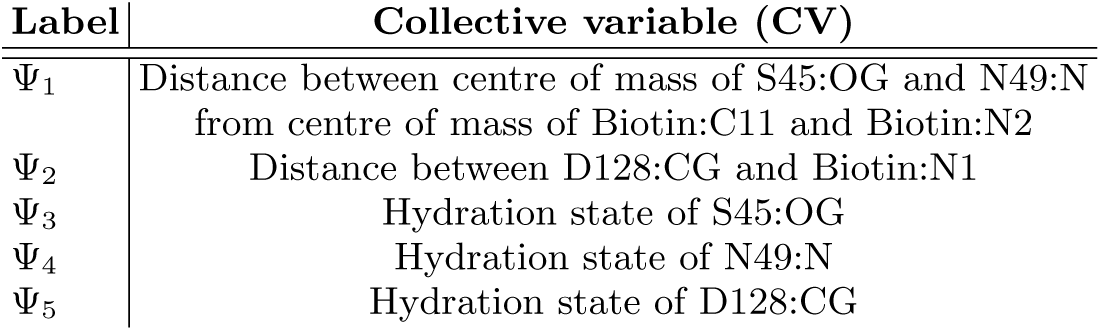
One-dimensional RC was constructed through SGOOP as a linear combination of the following 5 collective variables. See relevant residues marked in Fig. 4.

| Label | Collective variable (CV) |
| --- | --- |
| $\Psi_1$ | Distance between centre of mass of S45:OG and N49:N from centre of mass of Biotin:C11 and Biotin:N2 |
| $\Psi_2$ | Distance between D128:CG and Biotin:N1 |
| $\Psi_3$ | Hydration state of S45:OG |
| $\Psi_4$ | Hydration state of N49:N |
| $\Psi_5$ | Hydration state of D128:CG |

### Unbinding dynamics for WT and N23A can be mapped onto a one-dimensional reaction coordinate

Several groups over the decades have asked the question: does unbinding of biotin-streptavidin proceeds through well-defined ligand exit pathways that can be described by a reaction-coordinate (RC) model?^2,3^ For this system, the experimental evidence is in favor of this statement, given well-defined exponential statistics for the ligand residence times.^3^ Here, we explicitly construct such a RC using the method SGOOP, which allows optimizing a low-dimensional RC from a dictionary of possible collective variables or order parameters.^12^ The key idea in SGOOP is that the best RC will show a higher gap between the timescales associated with slow and fast processes, referred to in the literature as spectral gap (Methods). The dictionary of 5 such collective variables used for the systems here is detailed in Table 1, which comprises various protein-ligand contacts and hydration states of specific protein residues. For the dimeric biotin-streptavidin considered here, we find that both the WT and N23A unbinding can be reliably mapped into a 1d RC picture (Fig. 2) with very similar optimized RCs for WT and N23A. Fig. 2(a) shows the approximate 1-d free energy profile as a function of the trial RC used by SGOOP (see Methods) and as a function of the optimized RC. Fig. 2(b) shows that the spectral gap is indeed larger for the optimized RC. Corresponding results for N23A are given in supplementary results. Note that these free energies were generated using a sub-optimal RC and can not be expected to be fully converged. We have shown in the past that SGOOP works well such sub-optimal free energies.^18^

**FIG. 2:**
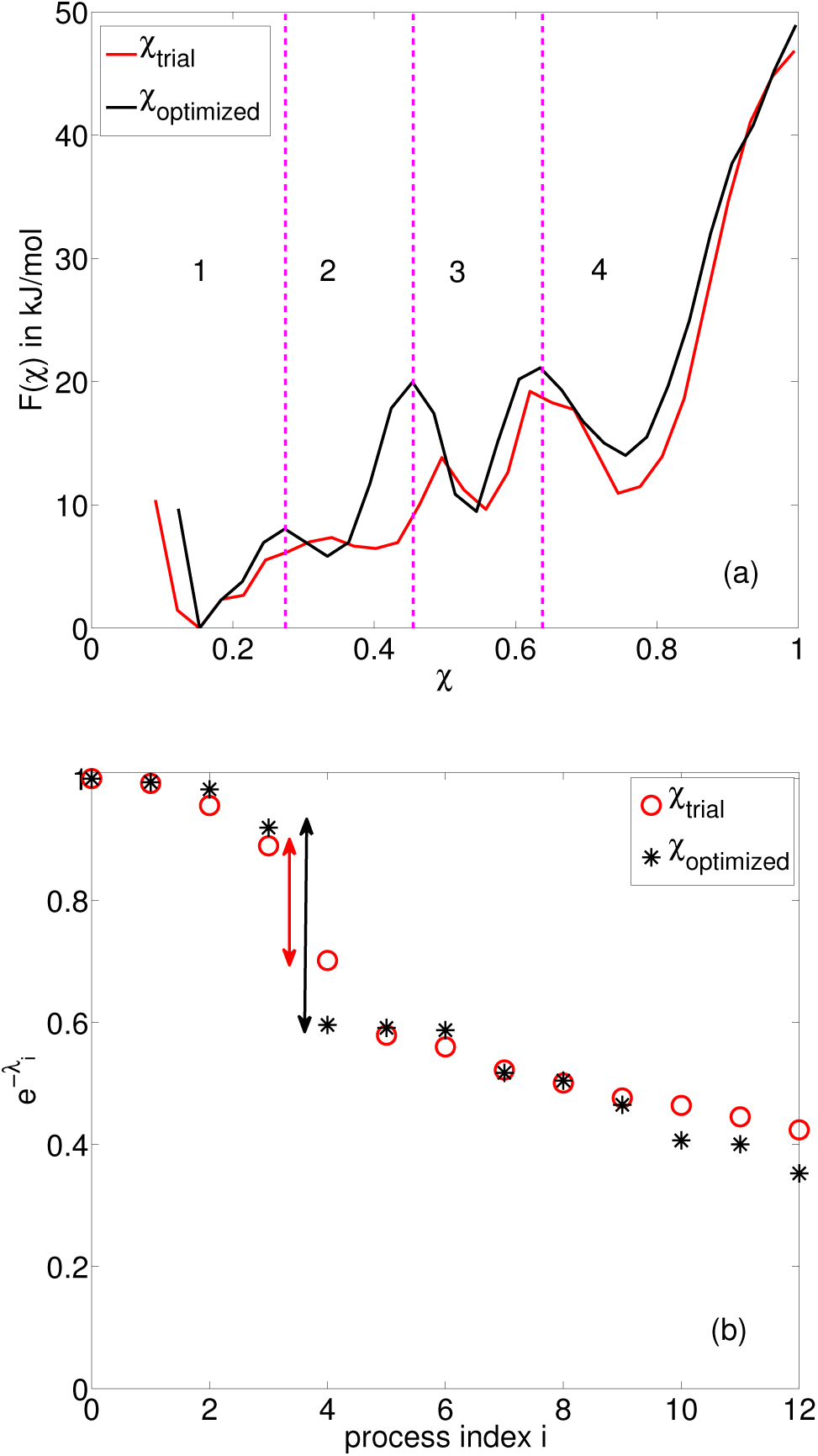
(a) Approximate free energies in kJ/mol as function of the trial and optimized RC (red and black respectively) for WT. To help with comparison, in this plot both RCs have been scaled by their respective maximum recorded values (reported in SR). 4 states are visible in this free energy profile, where state 1 is bound and state 4 is unbound. The other two are metastable intermediates. The very high free energy in the end of state 4 is because the runs were stopped when the trajectories reached state 4 and committed to it hence this state is not fully sampled. (b) Eigenvalue spectrum corresponding to trial and optimized RC (red circles and black asterisks respectively) for WT. Corresponding to the 3 barriers seen in (a) for the optimized RC, we mark the spectral gap as the difference between the eigenvalues for 4th and 5th eigenmodes (since the first eigenmode with *λ*_0_ = 0 is just the stationary state). It is clear how the SGOOP-identified RC has sharper and higher barriers in (a), and a much larger spectral gap in (b). Analogous plots for free energies and eigenvalue spectrum for the mutant N23A have been provided in the SR.

Another question which has received much attention concerns the nature of the coupling between protein hydration and the lengthening of a specific biotin hydrogen bonding interaction. Stayton et al and others have demonstrated through simulations that an early mechanistic event in biotin-streptavidin dissociation involves these two events (viz. D128 hydration and D128-biotin bond stretching) happening in a concerted manner.^2,3^ Here, we find through SGOOP that the optimized RC involves contribution from the D128-biotin bond stretching but not from the hydration state of D128 (Fig. 3), which is unequivocal evidence that, at least for this system, it is the D128-biotin bond stretching which is the bottleneck. Of course this picture will likely change for other systems - what is more important here is the demonstrated ability to answer such questions with quantitative confidence. In Supplementary Results, we give similar spectral gap plots for the full exploration of the 5-dimensional order parameter space for WT and N23A.

**FIG. 3:**
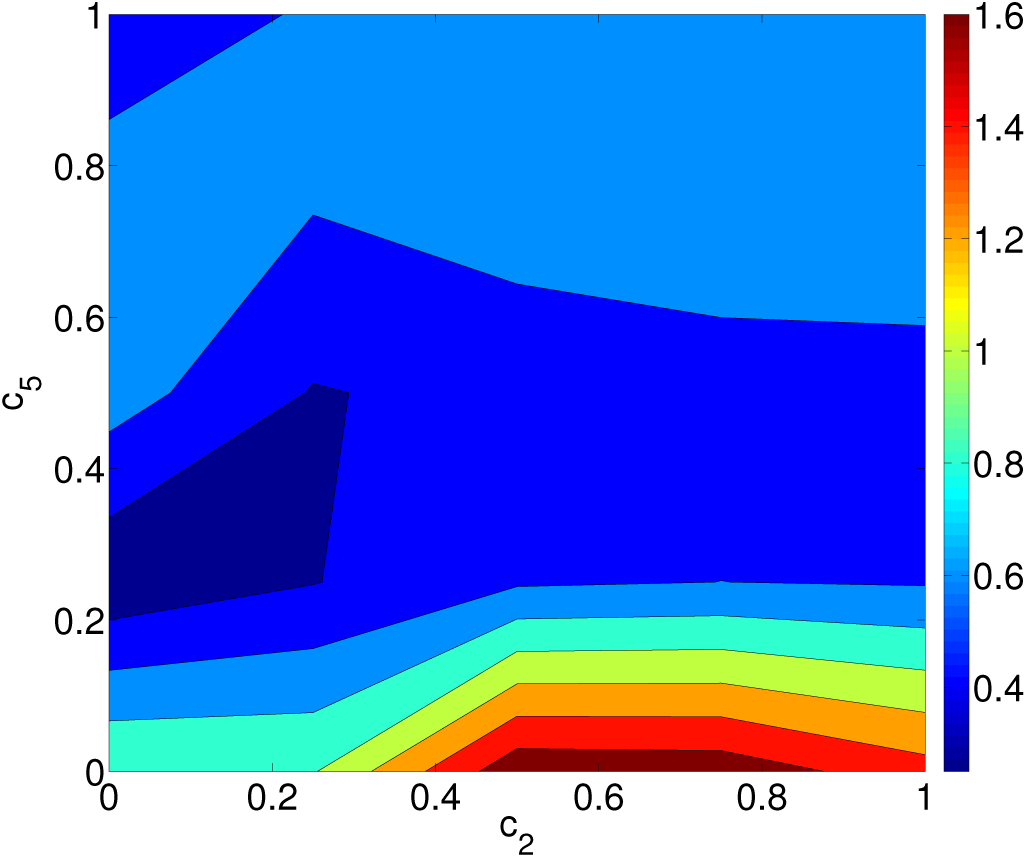
Here we show the relative contributions of D128-biotin bond stretching (*c*_2_) and D128 hydration (*c*_5_) to the spectral gap of the dynamics when viewed as function of different trial RC *χ* = *c*_1_Ψ_1_ + *c*_2_Ψ_2_ + *c*_5_Ψ_5_. The full dictionary {Ψ_*i*_} of collective variables is defined in Table 1. Similar spectral gap profiles for different 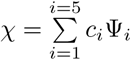 for WT and for N23A are provided in the SR. The spectral gap is in arbitrary units and has been scaled so that the case with *χ* ={1, 0, 0, 0, 0}, which was the trial RC, has a spectral gap of 1 unit. The optimized RC with highest spectral gap is *χ* = Ψ_1_ + 0.75Ψ_2_.

### Unbinding dynamics for WT and N23A can be mapped into 4 states and sequential movement between them

When viewed as a function of the optimized reaction coordinate, we find that the free energy landscape Fig. 2 for both the WT and N23A maps into 4 states, namely bound, unbound and 2 intermediates. Through infrequent metadynamics runs with the optimized RC (Methods) we find these to have diverse lifetimes across several order of magnitudes. Here we describe in details the states for the WT (Fig. 4 and Fig. S1 for WT and N23A respectively) - analogous description for N23A is given in the Supplementary Results (SR).

**FIG. 4:**
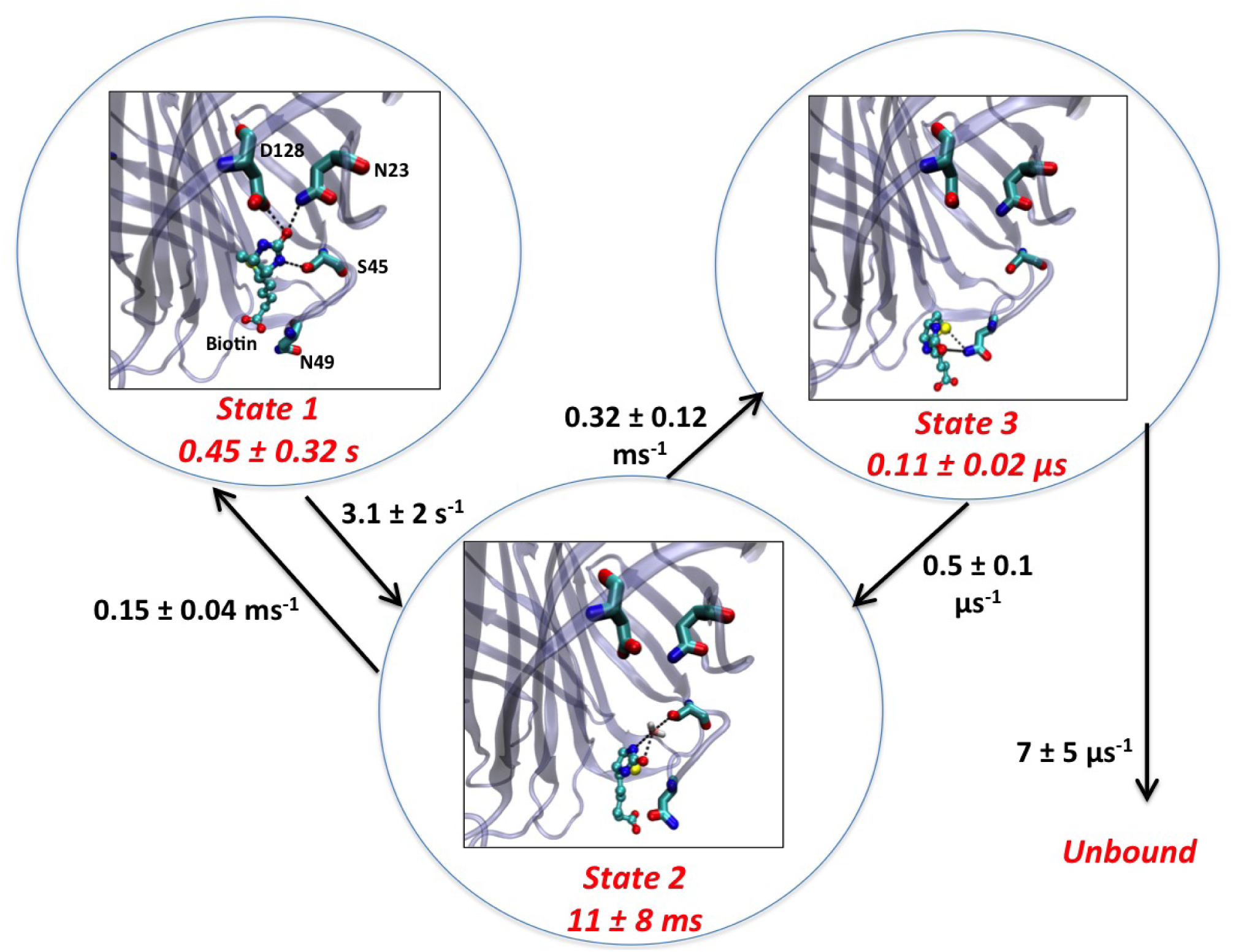
State-to-state transitions rates and various residence times for WT biotin-streptavidin dimer.

State 1 is the bound state with a lifetime of 0.45±0.32 sec. Some of the distinguishing and stabilizing interactions here include direct hydrogen bonds between biotin and the residues D128, N23 and S45. To leave state 1, the interactions of biotin with D128 and with N23 distort from being direct to water-mediated, and are finally broken. State 2 is a key intermediate state, with a lifetime of 11±8 msec. The characterizing interaction here is a water mediated bond between biotin and S45, and interactions between biotin and N49. Once the S45 interaction is broken, the system reaches state 3 which is the last intermediate state and has a lifetime of 0.1±0.02 *μ*sec. Here the biotin interacts mainly with N49 and adjoining residues on the periphery of the binding pocket. This state is composed of several shallow energy basins with fast interconversion. The biotin is almost fully solvent exposed now and finally it reaches state 4, which is the freely diffusing unbound state.

### N23A unbinding has same rate-determining step as WT, only 100 times faster

We extract the residence times in various states and rate constants for movements between them by pooling the various infrequent metadynamics runs (Methods). This gives a matrix of transition rates and a discrete-time master equation for movement between the various states. The eigenvalues and eigenvectors of this matrix carry precious information about the slow steps in unbinding and their time scales. We calculate this transition matrix and its eigenvalues for different values of minimum time the trajectory must spend in a certain state before being labelled as being *committed* to that state. Such a metric, introduced previously by Tiwary, Mondal and Berne^14^, is inspired by the notion of friction-induced spurious recrossing of the barrier between two state, and attempts to discount such recrossings from genuine transition events. This is akin to the lag time in markov models, and the dynamics can be trusted only for those eigenvalues which converge beyond a certain minimum commitment time. As can be seen in Figs. 5(a-b) the eigenvalues for both the WT and N23A are well-converged with respect to this metric.

**FIG. 5:**
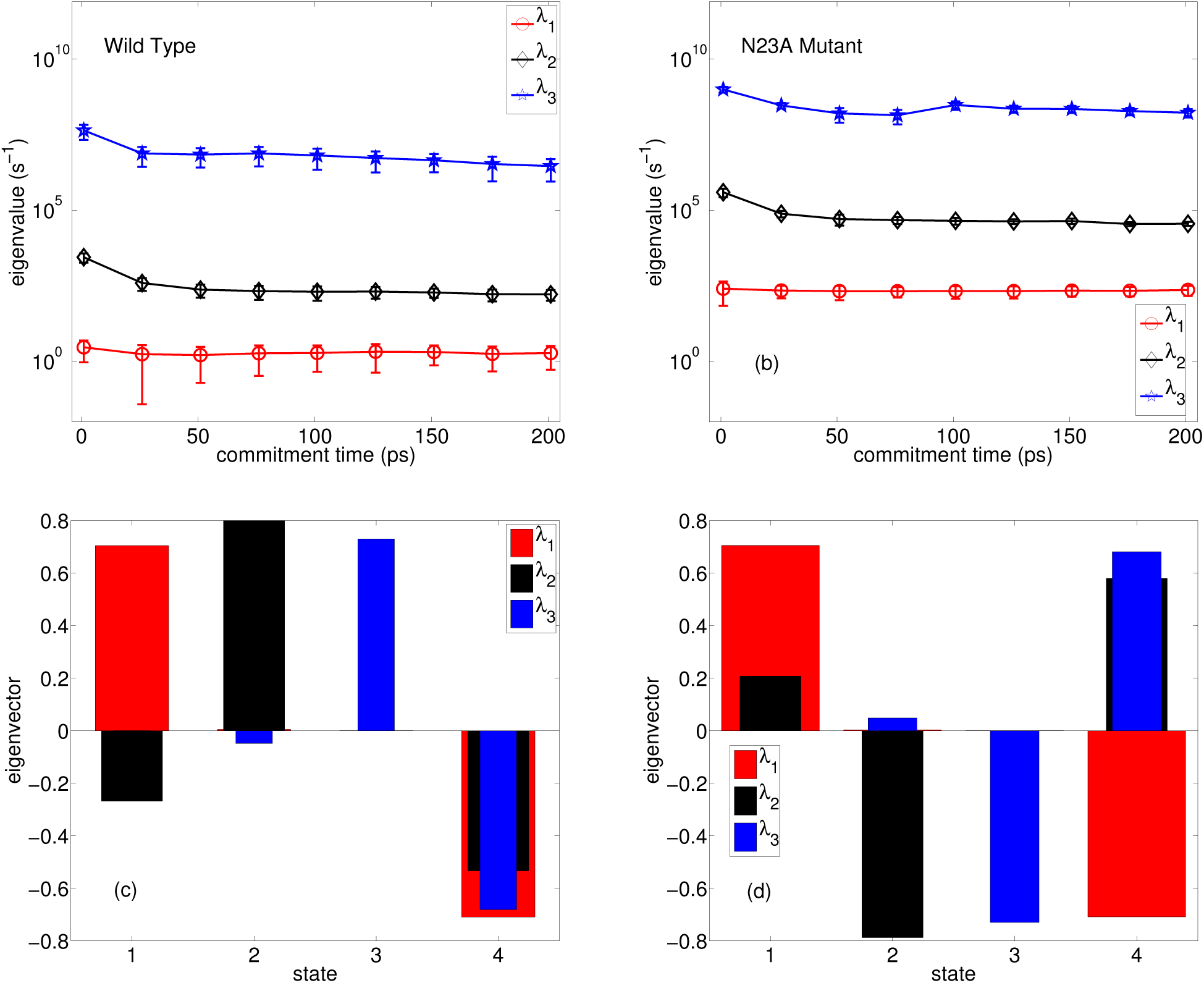
Top (a-b): Eigenvalues for the transition matrix for state-to-state movement for WT and N23A respectively. Red, blue and black denote slowest, second-slowest and third-slowest processes respectively. These are shown as a function of different minimum commitment time used to ascertain the convergence of these eigenvalues, as described in Results. Note how the slowest eigenvalue for N23A is 2 orders of magnitude faster than that for WT. Bottom (c-d) Eigenvectors for the various processes using the same color-coding, calculated using a commitment time of 40 ps.

Figs. 5(c-d) show the corresponding eigenvectors. Of special importance is the slowest eigenvalue and eigenvector - which corresponds to the dynamical bottleneck in unbinding of biotin-streptavidin. Interestingly, both for WT and N23A the exit from state 1 forms the bottleneck, as can be seen from the node in the eigenvector i.e. movement happens between states with maximally positive and negative eigenvectors. For N23A this step is around 100 times faster than for WT, which is explained by the missing N23-biotin interaction in the mutant. Once state 1 is exited, the rest of the unbinding process is similar for both forms.

## DISCUSSION

Biotin-streptavidin is a classic system used to gain insight into protein-ligand interactions. It is heavily used across the biotechnological and nanobiotechnological industry, and is well-known for its extremely long biotin residence times. In this work we studied the dimeric biotin-streptavidin system, which is slightly more tractable than the full tetrameric system, in its wild type and N23A mutated forms. Using new enhanced sampling methods^11,12^ we demonstrated that atomistic simulations could reproduce the experimentally reported variation in residence time due to this motivation. Our simulations go far beyond just being able to calculate residence times - we also identify the various molecular determinants and bottlenecks in the unbinding process and quantify their roles. One of these is the longstanding question regarding cause/effect in the coupled events of D128-biotin and bond stretching and water entry into the binding pocket to hydrate D128. We establish that at least in the current dimeric system (which itself has been the subject of previous investigations^21^), it is the bond stretching which forms the bottleneck in the early parts of the unbinding. Our reported residence times for the dimeric system are in rough agreement with what one expects after taking into account the difference in binding affinities between the dimeric and the tetrameric systems.

In this work we were able to map the dynamics on a one-dimensional RC as a combination of given collective variables. This is not entirely surprising given that the experimentally reported residence times for the tetrameric biotin-streptavidin system follow a mono-exponential distribution. However, in more complicated systems, there is no guarantee that a one-dimensional RC exists or can be found. We are however hopeful that the methods and statistical tests used in this work will signal in a self-consistent manner if either the dictionary of trial collective variables needs to be expanded, or if the one-dimensional RC framework is simply not sufficient. We have in fact found this to be the case for unbinding in still more complex systems such as some of those in Ref.^31^.

Encouraged by the success in this system, we are now in the process of looking at the full tetrameric system with a bigger range of mutations. One crucial difference between these two systems is that we anticipate the need to incorporate movement between the two dimeric units as an additional member of the dictionary of collective variables from which the RC is optimized. Even though Grubmüller *et al* comment in their original paper using a biotin-streptavidin monomer instead of tetramer^5^ that they do not expect this simplification to change the results, we wonder if we will see surprises when we simulate the full tetramer. Specifically, given the considerable protein fluctuations in specific surface loop regions (see Fig. 1), having the presence of a second dimer unit which clamps these fluctuations could have profound affects on the unbinding pathways. Furthermore, other systems involving biotin are now known to display considerable allosteric affects as well - often involving surface loops,^32,33^ and we wonder if we might find similar affects at work in the full tetramer. In summary, we expect that recent and continuing developments in atomistic simulations and enhanced sampling are now taking us into an exciting age of reliable and routine calculations of the dynamics of protein-ligand interactions.

## ACKNOWLEDGMENTS

The author is grateful to Dorothy Beckett for suggesting this system and for several helpful discussions, and to Bruce Berne and Michele Parrinello for countless inspiring discussions. The author acknowledges computational support from University of Maryland’s Deepthought cluster and from Extreme Science and Engineering Discovery Environment (XSEDE) [TG-MCA08X002], and financial support from University of Maryland start-up grant. The author also thanks Anthony Clark for help with system preparation, and Purushottam Dixit for explaining to him the principle of Maximum Caliber.

**Author contributions:** P.T. designed the research, performed calculations, analyzed data and wrote the manuscript. **Competing interests:** The authors declare that they have no competing interests.**Data and materials availability:** All data needed to evaluate the conclusions in the paper are present in the paper and/or the Supplementary Results. Additional data related to this paper may be requested from the author.

## METHODS

### SYSTEM SPECIFICATIONS

The initial structure of biotin bound to streptavidin in its dimeric form was obtained from protein data bank (PDB:3RY2). Amber ff99SB*-ILDN all-atom force-field and TIP4P water model were used.^34,35^ All the non-protein molecules were removed, and protein preparation wizard within Schrodinger’s software package Maestro was used to post process the structures of protein for simulation. The residue mutation N23A was also carried out using Maestro. Protonation states of the amino acid residues were assigned assuming the systems are at pH 7.0. The ligand charges were parametrized using the general amber forcefield and AM1-BCC charges using the software package Chimera.^36,37^ The protein and ligand structures were concatenated and then solvated with TIP4P water. Counterions in the form of sodium chloride (NaCl) salt were added at a concentration of 150 mM to maintain physiological conditions. The full systems, which comprised of 82,793 atoms for each of WT and N23A, including protein, solvent and charge-compensating counter-ions, were prepared and equilibrated. The full structure’s energy was minimized, and then further equilibration MD simulations was run first in the NVT (at temperature 300 K) and then in the NPT ensembles (at pressure 1 bar and temperature 300 K) for 1 ns each, keeping the position of ligand restrained in its binding pose. Nose Hoover Thermostat^38^ and Parinello-Rahman barostat^39^ were employed. Periodic boundary conditions were applied in all three directions. The equations of motion were evolved with a time step of 2 fs. All our subsequent simulations were performed using constant temperature and pressure ensemble (with isotropic pressure) at temperature *T* = 300 K using the software GROMACS 5.0.7 patched with the plugin PLUMED 2.1.^40,41^

### INFREQUENT METADYNAMICS FOR KINETICS

In this work, we use the infrequent metadynamics approach^11,30^ to obtain unbinding trajectories and kinetic rate constants. This involves periodic but infrequent biasing of a low-dimensional RC in order to increase the probability of escape from metastable states where the system would ordinarily be trapped for extended periods of time.^42^ If the RC can demarcate all relevant stable states of interest, and if the time interval between biasing events is infrequent compared to the transition path time spent in the ephemeral transition state (TS) regions, then one increases the likelihood of not adding bias in the TS regions and thereby keeping pristine the dynamics of barrier crossing. This in turn preserves the sequence of transitions between stable states that the unbiased trajectory would have taken.^11^ The acceleration *α* of transition rates through biasing which directly yields the true unbiased rates, is then through generalized transition state theory^11,43–45^:

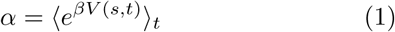

Here *s* is the RC being biased, *β* = 1*/k*_*B*_*T* is the inverse temperature, *V* (*s, t*) is the bias experienced at time *t* and the subscript *t* indicates averaging under the time-dependent potential. One can verify *a posteriori* if the requirements of Ref.^11,30^ were satisfied by checking if the cumulative distribution function for the transition times is Poisson^30^. Note that all rate constants are calculated through the use of the accelerated time^11,44^, without having to converge the free energy. Here we use SGOOP^12,25^ for the construction of a desirable RC (next section).

Total 5 collective variables (CVs) were used in this work (Table 1) and an optimized 1-dimensional RC *χ* was constructed as a linear combination of these 5 CVS using SGOOP (next section). As can be seen in Table 1, out of these 5 CVs 3 were distance variables *d* between the ligand and different parts of the protein. The other 3 were hydration CVs proportional to the number of water molecules in the vicinity of specific protein residues. Since the CVs need to be smoothly differentiable in order to be biased, the hydration variable *w* was implemented through the function

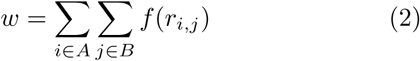

Here the set *A* denotes the respective protein residues from Table 1. The set *B* comprises all the oxygen atoms of various water molecules. *r*_*i,j*_ is the distance in *nm* between atoms *i* and *j* coming from these two sets respectively. The function

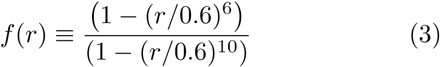

is a switching function that makes *w* smoothly differentiable.

As described in the following section and in Supplementary results, the optimized RC was found through SGOOP to be *χ* = *d*_1_ + 0.75*d*_3_. The bias was added once every 5 picoseconds as a Gaussian function of 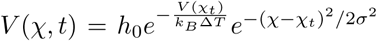. Here *χ*_*t*_ is the value of the RC at a certain time *t*, *h*_0_ is the initial Gaussian amplitude which we took as 2 kJ/mol. Δ*T*, which we took to be Δ*T* = 14 *T* is the so-called tempering parameter which smoothly modulates the amplitude of the Gaussian each time a certain point in the RC space is revisited.^42,46^ The Gaussian width *σ* was taken as 0.02 units. The free energy as a function of *χ*, defined below, can be estimated from the bias as (within an additive constant^47^):

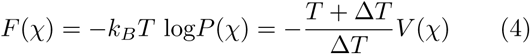

where *P* (*χ*) is the stationary probability density as a function of *χ*.

We define the unbound state as states with *χ* > 3, beyond which we find that the ligand diffuses in the bulk. All 14 independent simulations for both WT and N23A were ran until they gave full unbinding. This took between 2 nanosecond of unscaled MD time for the fastest exit event to as much as 11 nanoseconds for the slowest event. Through each of the simulations we found no deterioration of the protein structure as measured by the RMSD value of the heavy atoms, which stayed within 3 Å of the native pose (see Supplementary Results for RMSD plots).

### SGOOP FOR REACTION COORDINATE OPTIMIZATION

Spectral gap optimization of order parameters (SGOOP)^12,25^ is a method to optimize a RC as a combination (linear or non-linear) of a set of many given candidate collective variables Ψ = (Ψ_1_, Ψ_2_, …, Ψ_*d*_). SGOOP considers the optimal RC to have the following characteristics: (1) it can demarcate between various metastable states that the full system possesses, and (2) it can maximize separation of timescales between visible slow and hidden fast processes. This timescale separation is calculated as the spectral gap between the slow and fast eigenvalues of the transition probability matrix on a grid along a trial RC^12,25^. The transition probability matrix is calculated in SGOOP using an approximate kinetic model that can be derived for example through the principle of Maximum Caliber.^23–25^ Let {*λ*} denote the set of eigenvalues of this matrix, with *λ*_0_ = 0 < *λ*_1_ ≤ *λ*_2_ ≤ …. We refer to 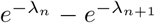 as the spectral gap, where *n* is the number of barriers apparent from the free energy estimate that are higher than a user-defined threshold (typically ≳ *k*_*B*_*T*). Note that this calculation is done without any pre-knowledge of the number of metastable states the system might have, and as such makes it easy to use SGOOP for systems with an arbitrary number of metastable states as in this work.

The key input to SGOOP is an estimate of the stationary probability density *P* (*χ*) (or equivalently the free energy) of the system, accumulated through a biased simulation performed along a sub-optimal trial RC *χ* given by some linear or non-linear function *χ* = *f*_0_(Ψ), where Ψ denotes the larger set of candidate CVs. This is done by performing 8 independent *frequent* metadynamics runs (biasing interval = 2 ps)^42,47^ and using Eq. 4 to average over the estimate from these runs. The bias deposition rate is higher for these preliminary runs since the objective in this part is to sample the free energy landscape and not necessarily preserve kinetic pathways. Metadynamics^47^ allows to easily reweight for the stationary probability density for any RC in post-processing without having to repeat simulations. This stationary density estimate is complemented with short unbiased MD runs (10 ns long) in the bound pose, which provide the dynamical observables for SGOOP, namely the average number of transitions per unit time along any trial RC. These serve as a proxy for the explicit knowledge of the diffusivity tensor. Given these pieces of information we use the principle of Maximum Caliber^23–25^ to set up an unbiased master equation for the dynamics of various trial CVs *f* (Ψ). Through a post-processing optimization procedure we then find the optimal RC as the *f* (Ψ) which gives the maximal spectral gap of the associated transfer matrix, and hence the optimized RC. We refer to Ref. 25 and to Supplementary Results for further details of the theory behind SGOOP and its application in this work.

## References

1. P. C. Weber, D. Ohlendorf, J. Wendoloski, and F. Salemme, Science 243, 85 (1989).

2. D. E. Hyre, L. M. Amon, J. E. Penzotti, I. Le Trong, R. E. Stenkamp, T. P. Lybrand, and P. S. Stayton, Nat. Struc. Mol. Bio. 9, 582 (2002).

3. S. Freitag, V. Chu, J. E. Penzotti, L. A. Klumb, R. To, D. Hyre, I. Le Trong, T. P. Lybrand, R. E. Stenkamp, and P. S. Stayton, Proc. Natl. Acad. Sci. 96, 8384 (1999).

4. K. Caswell, J. N. Wilson, U. H. Bunz, and C. J. Murphy, Journal of the American Chemical Society 125, 13914 (2003).

5. H. Grubmüller, B. Heymann, and P. Tavan, Science, 997 (1996).

6. T. Young, R. Abel, B. Kim, B. J. Berne, and R. A. Friesner, Proceedings of the National Academy of Sciences 104, 808 (2007).

7. A. Chilkoti and P. S. Stayton, Journal of the American Chemical Society 117, 10622 (1995).

8. R. A. Copeland, Nat. Rev. Drug. Discov. 15, 87 (2016).

9. C. S. Tautermann, Curr. Opin. Pharm. 30, 22 (2016).

10. A. Dickson, P. Tiwary, and H. Vashisth, Current topics in medicinal chemistry (2017).

11. P. Tiwary and M. Parrinello, Phys. Rev. Lett. 111, 230602 (2013).

12. P. Tiwary and B. J. Berne, Proc. Natl. Acad. Sci. 113, 2839 (2016).

13. P. Tiwary, V. Limongelli, M. Salvalaglio, and M. Parrinello, Proc. Natl. Acad. Sci. 112, E386 (2015).

14. P. Tiwary, J. Mondal, and B. J. Berne, Science Advances 3 (2017).

15. R. Casasnovas, V. Limongelli, P. Tiwary, P. Carloni, and M. Parrinello, Journal of the American Chemical Society 139, 4780 (2017).

16. Y. Wang, E. Papaleo, and K. Lindorff-Larsen, eLife 5, e17505 (2016).

17. J. I. Stuckey, B. M. Dickson, N. Cheng, Y. Liu, J. L. Norris, S. H. Cholensky, W. Tempel, S. Qin, K. G. Huber, C. Sagum, et al., Nature chemical biology 12, 180 (2016).

18. P. Tiwary and B. J. Berne, J. Chem. Phys. 145, 054113 (2016).

19. K. L. Fleming, P. Tiwary, and J. Pfaendtner, The Journal of Physical Chemistry A 120, 299 (2016).

20. D. Bochicchio, M. Salvalaglio, and G. M. Pavan, Nature Communications 8 (2017).

21. F. M. Aslan, Y. Yu, S. Vajda, S. C. Mohr, and C. R. Cantor, Journal of biotechnology 128, 213 (2007).

22. L. A. Klumb, V. Chu, and P. S. Stayton, Biochemistry 37, 7657 (1998).

23. S. Pressé, K. Ghosh, J. Lee, and K. A. Dill, Rev. Mod. Phys. 85, 1115 (2013).

24. P. D. Dixit, A. Jain, G. Stock, and K. A. Dill, J. Chem. Theor. Comp. 11, 5464 (2015).

25. P. Tiwary and B. J. Berne, The Journal of Chemical Physics 147, 152701 (2017).

26. N. Agmon and J. Hopfield, The Journal of chemical physics 79, 2042 (1983).

27. G. G. Poon and B. Peters, The Journal of Physical Chemistry B 120, 1679 (2015).

28. C. Moritz, A. Trster, and C. Dellago, The Journal of Chemical Physics 147, 152714 (2017).

29. A. Berezhkovskii and A. Szabo, The Journal of chemical physics 122, 014503 (2005).

30. M. Salvalaglio, P. Tiwary, and M. Parrinello, J. Chem. Theor. Comp. 10, 1420 (2014).

31. F. E. Kwarcinski, K. R. Brandvold, S. Phadke, O. M. Beleh, T. K. Johnson, J. L. Meagher, M. A. Seeliger, J. A. Stuckey, and M. B. Soellner, ACS Chem. Bio. 11, 1296 (2016).

32. L. H. Weaver, K. Kwon, D. Beckett, and B. W. Matthews, Proceedings of the National Academy of Sciences 98, 6045 (2001).

33. E. D. Streaker and D. Beckett, Journal of molecular biology 292, 619 (1999).

34. K. Lindorff-Larsen, S. Piana, K. Palmo, P. Maragakis, J. L. Klepeis, R. O. Dror, and D. E. Shaw, Proteins 78, 1950 (2010).

35. W. L. Jorgensen, J. Chandrasekhar, J. D. Madura, R. W. Impey, and M. L. Klein, J. Chem. Phys. 79, 926 (1983).

36. J. Wang, R. M. Wolf, J. W. Caldwell, P. A. Kollman, and D. A. Case, Journal of computational chemistry 25, 1157 (2004).

37. E. F. Pettersen, T. D. Goddard, C. C. Huang, G. S. Couch, D. M. Greenblatt, E. C. Meng, and T. E. Ferrin, Journal of computational chemistry 25, 1605 (2004).

38. D. J. Evans and B. L. Holian, The Journal of chemical physics 83, 4069 (1985).

39. M. Parrinello and A. Rahman, Physical Review Letters 45, 1196 (1980).

40. B. Hess, C. Kutzner, D. Van Der Spoel, and E. Lindahl, J. Chem. Theor. Comp. 4, 435 (2008).

41. G. A. Tribello, M. Bonomi, D. Branduardi, C. Camilloni, and G. Bussi, Comp. Phys. Comm. 185, 604 (2014).

42. O. Valsson, P. Tiwary, and M. Parrinello, Ann. Rev. Phys. Chem. 67, 159 (2016).

43. B. J. Berne, M. Borkovec, and J. E. Straub, J. Phys. Chem. 92, 3711 (1988).

44. A. F. Voter, Phys. Rev. Lett. 78, 3908 (1997).

45. H. Grubmüller, Phys. Rev. E 52, 2893 (1995).

46. A. Barducci, G. Bussi, and M. Parrinello, Phys Rev Lett 100, 020603 (2008).

47. P. Tiwary and M. Parrinello, J. Phys. Chem. B 119, 736 (2014).

